# An inducible tricolor reporter mouse for simultaneous imaging of lysosomes, mitochondria and microtubules

**DOI:** 10.1101/2023.05.22.541817

**Authors:** Vera Hutchison, Anne Lynch, Andrés Mauricio Gutierrez Gamez, Jichao Chen

## Abstract

Cell-type-specific use of the same DNA blueprint generates diverse cell types. Such diversity must also be executed via differential deployment of the same subcellular machinery. However, our understanding of the size, distribution, and dynamics of subcellular machinery in native tissues, and their connection to cellular diversity, remain limited. We generate and characterize an inducible tricolor reporter mouse, dubbed “kaleidoscope”, for simultaneous imaging of lysosomes, mitochondria and microtubules in any cell type and at a single cell resolution. The expected subcellular compartments are labeled in culture and in tissues with no impact on cellular and organismal viability. Quantitative and live imaging of the tricolor reporter captures cell-type-specific organelle features and kinetics in the lung, as well as their changes after Sendai virus infection. *Yap/Taz* mutant lung epithelial cells undergo accelerated lamellar body maturation, a subcellular manifestation of their molecular defects. A comprehensive toolbox of reporters for all subcellular structures is expected to transform our understanding of cell biology in tissues.

**SUMMARY STATEMENT:** Our understanding of subcellular machinery is often inferred from that in cultured cells. Hutchison et al. have generated a tricolor tunable reporter mouse for simultaneous imaging of lysosomes, mitochondria and microtubules in the native tissues at a single-cell resolution.

## INTRODUCTION

Nearly all cells in our body have the same DNA and subcellular machinery, with few exceptions due to specialization such as somatic mutations for adaptive immunity and denucleation of red blood cells. Selective use of the same DNA blueprint underlies our ∼200 cell types and must be executed via cell-type-specific use of the subcellular machinery, such as the T-tubules of muscles, microvilli of absorbent gut cells, synaptic junctions of neurons, and lysosomal related organelles of melanocytes. Besides such causation, a cell type is associated with and can be defined by not only expressed genes, but also cellular features, as exemplified in Ramon y Cajal’s neuronal classification by cell morphology^1^. Moreover, the myriad DNA-coded macromolecules, as well as their cofactors, substrates and products, are confined to organelles, transported via the vesicular system along the cytoskeleton, and connected through cell-cell and cell-matrix adhesions.

Despite this analogy and interplay between molecular and cellular features, the former has enjoyed robust understanding and tools, leading to a conceptual framework where combinatorial action of transcription factors drives cell-type-specific epigenomic changes and consequently gene expression. In contrast, studies of cellular features are typically limited to cultured cells due to technical feasibility. However, as genes have distinct functions across cell types, subcellular machinery may also function differently in culture versus in tissues, particularly between large thin cells on 2D stiff plastics and those embedded in 3D cellular and matrix environment. Although classic transmission electron microscopy can pinpoint many subcellular structures, it is challenging to assign them to specific cell types, track them over time, and construct a 3D view for the entire cell.

The necessity to study cellular features in tissues and the difficulty in doing so are accentuated in the gas-exchanging lung alveoli, where a multitude of cell types interdigitate within thin tissues bordering millions of air pockets. For example, individual alveolar type 1 (AT1) cells and Cap2 endothelial cells (also known as Car4 cells or aCaps) span several hundred microns and multiple alveoli, whereas mesenchymal cells are as diverse in morphology and location^2–6^. Furthermore, their cellular processes are often too thin and intertwined to attribute subcellular staining signals to their cellular source. Inspired by the sparse cell labeling strategy achieved originally via Golgi staining and nowadays by genetic recombination^7^, we have visualized the diverse morphology of each cell type in the alveolar epithelial, endothelial, and mesenchymal lineages using a Cre-dependent cell membrane or cytosolic reporter^2, 5, 6^.

Expanding this pipeline of cell biology analysis, our current study has generated a reporter mouse to fluorescently label subcellular structures including lysosomes, mitochondria, and microtubules. Unlike existing reporters that are limited to neurons or individual structures^8–13^, our reporter enables tricolor labeling within a single cell of any type for which a Cre driver is available. Applying this reporter across organs and cell lineages, using live imaging, in a viral injury model, and in conjunction with genetic mutations, we have shown in native tissues cell-type-specific distributions of lysosomes and mitochondria, lysosomal fusion and fission, organelle changes upon viral infection, and lamellar body biogenesis and its defects in *Yap/Taz* mutant alveolar type 2 (AT2) cells.

## RESULTS

### Concatenated fluorescent fusion proteins label three subcellular structures when transiently expressed

Recognizing the use of *ROSA* reporters in not only lineage tracing, but also visualizing cell morphology (*ROSA^mTmG^* and *ROSA^tdT^*) and purifying ribosomes (*ROSA^L10GFP^*) or nuclei (*ROSA^Sun1GFP^*)^14–17^, we sought to systematically label subcellular structures, leveraging myriad fluorescent fusion proteins verified in cultured cells, to fill the gap in our knowledge of cell biology in the native tissues. To achieve comparable expression among multiple fusion proteins, we concatenated them with 2A self-cleaving sequences, instead of IRES^18–21^, and positioned the largest and thus potentially less robust (tag)BFP-Lamp1 in the preferred first position, before a mitochondrial localization signal (MLS)-linked mKate2 (a tagRFP variant) and an EGFP-tagged alpha-Tubulin (Fig. 1A, S1A). We named this tri-fusion protein construct “kaleidoscope” given its ability of multicolor labeling of structures inside a cell. We placed the kaleidoscope construct under the control of a strong ubiquitous CAG promoter used in other ROSA reporters, separated by a Cre-dependent transcriptional stop cassette.

**Figure 1:**
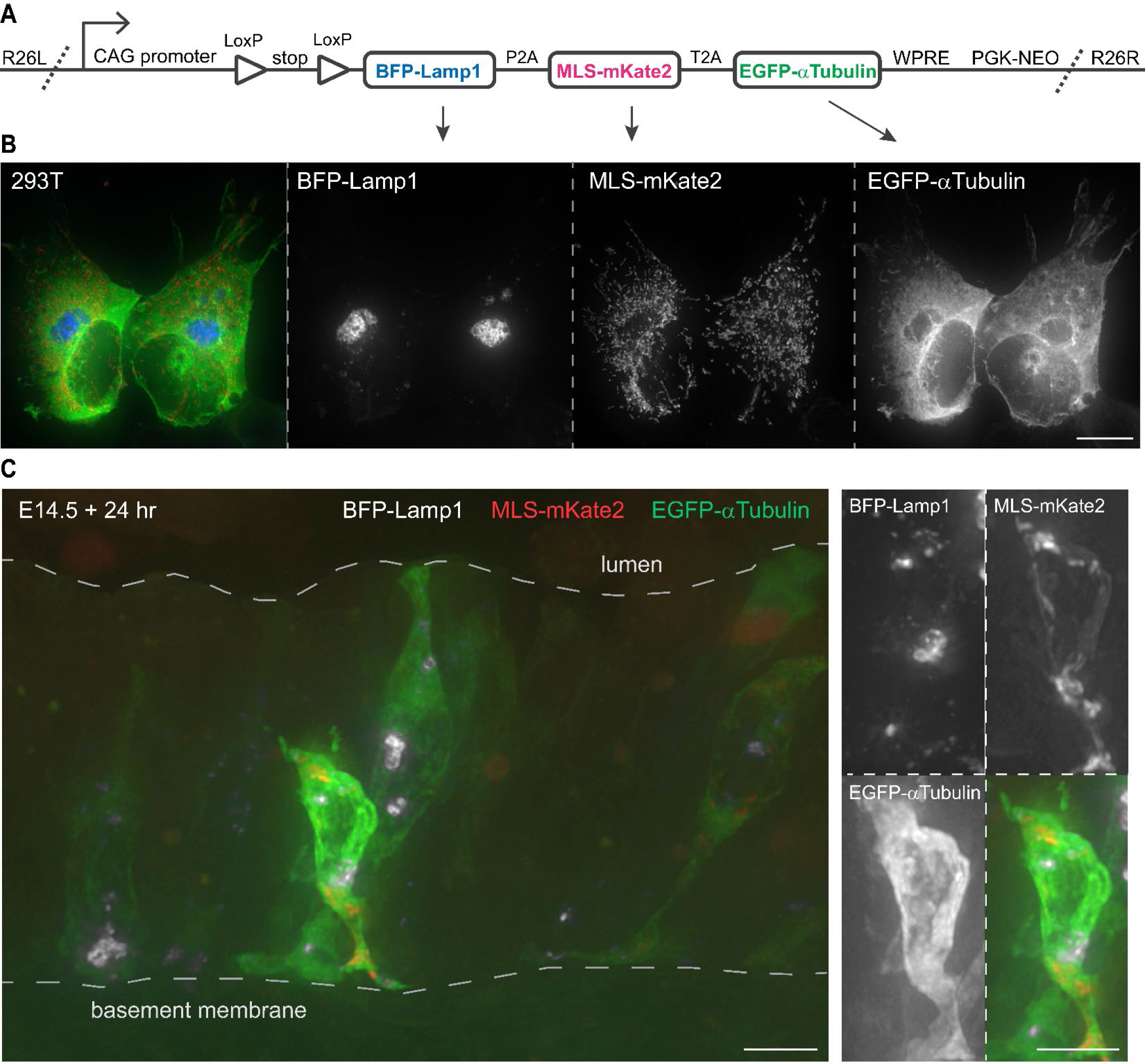
Design and in vitro validation of a Kaleidoscope reporter. (**A**) The Kaleidoscope reporter construct diagram showing 3 fluorescent fusion proteins connected with 2A self-cleaving sequences (P2A and T2A) driven by a strong, constitutive promoter (CAG), interrupted by a transcriptional stop flanked by LoxP sites, followed by a stabilizing Woodchuck hepatitis virus posttranscriptional regulatory element (WPRE) and a positive selection cassette (PGK-NEO), and sandwiched between left and right homology arms targeting the ROSA26 locus (R26L and R26R). (**B**) Representative images of HEK 293T cells co-transfected with ROSA-Kaleidoscope and pCAG-Cre, showing distinct subcellular localization of the 3 fusion proteins, color-coded as in panel A. Scale: 10 um. (**C**) Sporadic epithelial cells in an E14.5 mouse lung electroporated with Cre-recombined ROSA-Kaleidoscope and cultured for 24 hr. The enlarged cell is truncated by physical sectioning and thus does not reach the airway lumen, as the dimmer cell behind it does. Scale: 10 um.

To test the kaleidoscope construct, we removed the transcriptional stop cassette in vitro with a Cre recombinase and transiently transfected the Cre-recombined construct into a HEK293T human cell line^22^. Native fluorescence from BFP-Lamp1 exhibited a few perinuclear globules, MLS-mKate2 a vesicular network, and EGFP-alpha-Tubulin a reticular pattern excluding the cell nucleus and BFP-Lamp1 globules – distributions consistent with lysosomes, mitochondria, and microtubules, respectively (Fig. 1B). A similar pattern was observed in a mouse lung epithelial cell line MLE15^23^ (Fig. S1B). To further examine the construct in tissues, we electroporated it into embryonic mouse lungs and, after 24 hr culture, observed individual epithelial cells bridging the basement membrane and apical lumen and fluorescing from the 3 fusion proteins with expected distinct distributions (Fig. 1C).

### A tricolor reporter mouse labels lysosomes, mitochondria, and microtubules across organs

After validating the kaleidoscope fusion proteins via transient expression in cultured cell lines and lungs, we knocked the construct into the *ROSA* locus via CRISPR-stimulated homologous recombination in mouse embryonic stem cells (Fig. S1C). To examine the fusion proteins across organs, we activated the resulting *ROSA^Kaleidoscope^* allele with a near ubiquitous Cre driver, CMV-Cre^24^. As cell spreading on a 2D culture surface facilitates resolution of subcellular structures, we grew primary cells from dissociated postnatal day 7 (P7) lungs in conditions that most likely favor fibroblasts^25^ and compared the kaleidoscope fusion proteins with established markers of lysosomes (LAMP2), mitochondria (TOMM22), and microtubules (beta-Tubulin), with each pair showing remarkable overlaps (Fig. 2A). The EGFP-alpha-Tubulin was not as filamentous as beta-Tubulin despite the common use of this fusion protein^26, 27^, possibly due to extra fusion proteins unincorporated into the microtubule polymers. This was further examined later in the context of cell division. Western blots of whole lungs, in comparison with HEK293T cells transfected with individual fusion proteins, showed consistent patterns except for a lung-specific band possibly due to differential glycosylation of LAMP1 in tissues (Fig. S1D).

**Figure 2:**
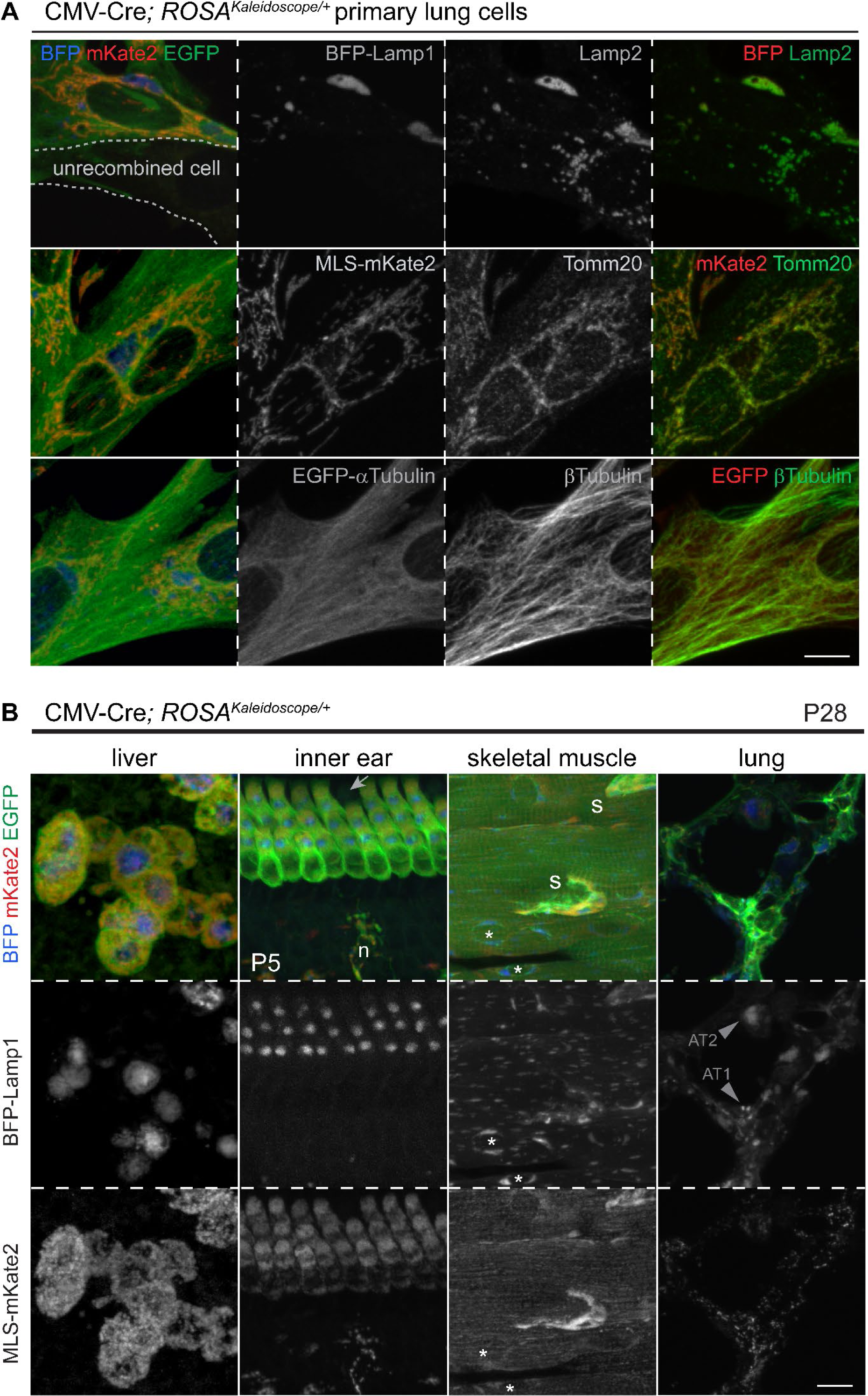
*ROSA^Kaleidoscope^* mice express correctly localized fusion proteins across cell types. (**A**) Primary cell culture from dissociated P7 CMV-Cre; *ROSA^Kaleidoscope/+^* lungs, showing colocalization of each fusion protein with markers of their expected subcellular structures: Lamp2 for lysosomes, Tomm20 for mitochondria, beta-Tubulin for microtubules. An escaper of CMV-Cre recombination is outlined with dashes. Scale: 10 um. (**B**) A survey of cell types and tissues from CMV-Cre; *ROSA^Kaleidoscope/+^* mice at P28, except for the inner ear at P5 to avoid calcification). Lysosomes range in size from 10 to 1 um characteristic of cell types in the decreasing order of hepatocytes, lung alveolar type 2 (AT2) cells, hair cells, skeletal muscle cells, and lung alveolar type 1 (AT1) cells. In the skeletal muscle, microtubules perpendicularly bundle actomyosin fibers, whereas lysosomes and mitochondria align with and fit between actomyosin fibers. Additional lysosomes surround the nuclei (asterisk). Arrow, unrecombined hair cell; n, neuronal cell; s, satellite cell. Scale: 10 um.

The CMV-Cre; *ROSA^Kaleidoscope^* mice loaded with multiple fusion proteins were healthy and fertile, and showed typical multi-nucleation in the liver, striation in the skeletal muscles, and a regular array of hair cells in the inner ear (Fig. 2B). Lysosomes were consistently perinuclear but differed in size characteristic for each cell type including, from the largest to the smallest, hepatocytes, outer hair cells, and skeletal muscle cells (Fig. 2B, S2A, S2B). Interestingly, microtubules were aligned perpendicular to skeletal muscle fibers, likely to organize actomyosin bundles, whereas lysosomes and mitochondria showed a longitudinal pattern in the remaining space (Fig. 2B, S2A). Unlike organs dominated by a major cell type, ubiquitous labeling in the lung was uninformative, as its alveolar region has dozens of cell types with intertwined cellular processes (Fig. 2B) – a major rationale to build this genetic reporter for sparse, cell-type-specific labeling, as illustrated below. Nevertheless, the airway epithelium was relatively simple and labeled cells were readily recognizable as ciliated cells with apical microtubule clusters and club cells with apical domes (Fig. S2C, S2D). Comparing across organs, cell types, and ages, we noticed tissue autofluorescence from elastin fibers in the adult lung obscured the kaleidoscope EGFP fusion protein, requiring amplification with a GFP antibody. Relatedly, mKate2 was readily detected in the Cy3 channel, sparing the Cy5 channel for antibody or nuclei staining.

### Visualization of microtubule dynamics through cell cycle phases

To further validate the microtubule labeling that was less discrete than the two organelle labelings, we reasoned that mitotic spindle assembly during mitosis would be an informative test case. Reassuringly, in cultured primary lung cells from our reporter mice, the filamentous microtubule network of interphase cells organized into the classic metaphase starbursts in an elevated, rounded cell body (Fig. 3A). To track microtubule dynamics in tissues, we used *S*ftpcCreER ^28^ to activate the kaleidoscope reporter in neonatal alveolar type 2 (AT2) cells with ongoing developmental proliferation. We identified microtubule labeling patterns consistent with textbook examples of interphase, prophase, metaphase, anaphase, and telophase, where EGFP-alpha-Tubulin cycled through configurations of a diffuse network, paired foci, paired starbursts, and a bridging bundle (Fig. 3B). Interestingly, while the labeled mitochondria reticulum was largely evenly split between the two separating daughter cells, a single dominant lysosome sometimes formed during mitosis, suggesting delayed assembly or synthesis of new lysosomes that was nevertheless compatible with cell division (Fig. 3B).

**Figure 3:**
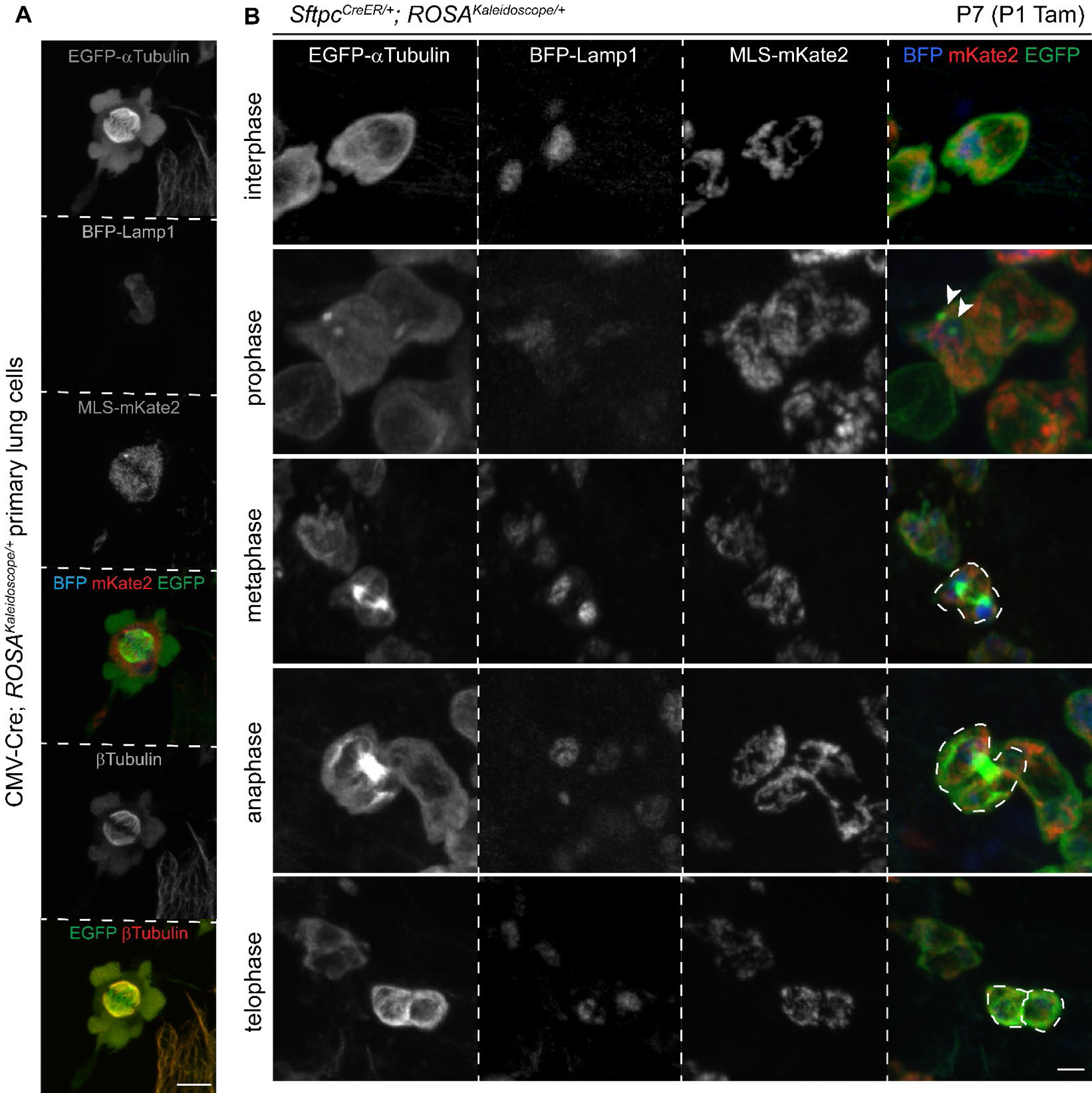
*ROSA^Kaleidoscope^* captures stereotypic mitotic microtubule structures. (**A**) Primary cell culture from dissociated P7 CMV-Cre; *ROSA^Kaleidoscope/+^* lungs with a mitotic spindle labeled with the EGFP-alpha-Tubulin fusion protein and colocalized with beta-Tubulin immunostaining, in contrast to their distribution in the adjacent interphase cell. Scale: 5 um. (**B**) Genetically labeled AT2 cells showing the distributions of EGFP-alpha-Tubulin fusion protein through the cell cycle stages. Arrowhead, centrosomal microtubules. Dividing daughter cells are outlined with dashes. Mitochondria fill both daughter cells, whereas lysosomes could be asymmetric. Tam, 100 ug tamoxifen. Scale: 5 um.

### Sizes and distributions of lysosomes, mitochondria and microtubules in AT2, AT1 and Cap2 cells

Unlike the small, conventionally shaped AT2 cells, alveolar type 1 (AT1) cells and Cap2 endothelial cells span several hundred microns and contribute to multiple alveoli and capillary segments, respectively^2, 5^. More challengingly, both large cells have thin cytoplasm at or below the resolution of optical microscopy and juxtaposed with cytoplasm from other cells, precluding attempts to assign a standard subcellular staining to a given cell type. Accordingly, we used *Rtkn2^CreER^* ^29^ and newly generated *Car4^CreER^* (Fig. S3A, S3B) to activate the kaleidoscope reporter in sporadic AT1 and Cap2 cells, respectively, with a limiting dose of tamoxifen – a sparse labeling strategy deployed to distinguish adjacent cells of the same type.

This revealed for the first time the complete distributions of lysosomes and mitochondria in AT1 and Cap2 cells in the native tissue (Fig. 4). Quantitative imaging showed that AT2 cells had one or two lysosome globules, likely corresponding to lamellar body clusters (verified below with LAMP3 immunostaining) and a rich mitochondrial network, which were both localized to the perinuclear region and respectively occupied around 5 and 15% of the cell volume defined by EGFP-alpha-Tubulin. AT1 cells had close to 50 lysosomal and 100 mitochondrial objects that were smaller or more fragmented and occupied a smaller fraction of the cell volume than those of AT2 cells. Both organelles were distributed throughout the expansive cytoplasm, forming mitochondrial and lysosomal outposts presumably to support active cellular metabolism distant from the nucleus (Fig. 4A, Supplementary Video 1). Cap2 cells had fewer but still abundant lysosomes and mitochondria that tended to accumulate near the nucleus as well as along the microtubule fibers, but infrequently at the cell extremities (Fig. 4A, Supplementary Video 2). The cuboidal AT2 cells were filled with a dense microtubule network, whereas AT1 cells had a mesh of filamentous microtubules, potentially providing mechanical support for their expansive and thin cytoplasm (Fig. 4A). By comparison, Cap2 cells had multiple long microtubule fibers, possibly due to their complex autocellular and intercellular junctions in the 3D capillary tubes (Fig. 4A). The differences in the microtubule density among cell types were reminiscent of those among a cotton ball (AT2 cells), a sieve (AT1 cells), and a fishnet (Cap2 cells) (Fig. S3C).

**Figure 4:**
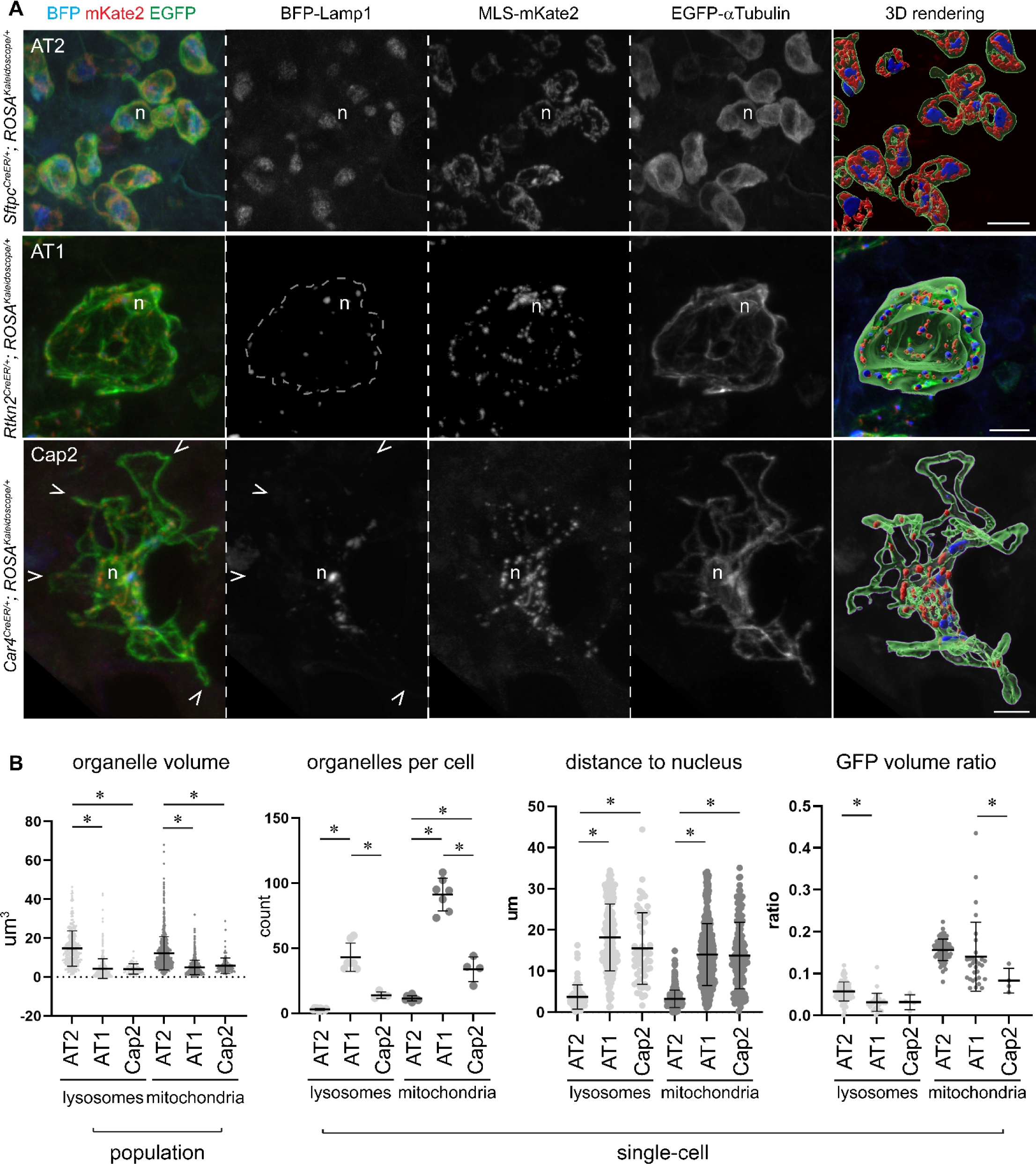
Cell-type-specific sizes and distributions of lysosomes and mitochondria in the lung (**A**) Genetic labeling and Imaris 3D rendering of lysosomes, mitochondria, and microtubules in individual AT2 (*Sftpc*^*CreER*^; 100 ug tamoxifen at P1 and harvested at P7), AT1 (*Rtkn2^CreER^*; 50 ug tamoxifen at P1 and harvested at P11), Cap2 (*Car4^CreER^*; 100 ug tamoxifen at P3 and harvested at P12) cells. n, nucleus of labeled cells based on exclusion of EGFP-alpha-Tubulin; dash, AT1 cell boundary; caret, Cap2 cell edge with few lysosomes and mitochondria. Scale: 10 um. See also Supplementary Video 1, 2. (**B**) Population and single-cell level measurements with means and standard deviations of lysosomes and mitochondria in genetically labelled AT2, AT1, Cap2 cells. Individual lysosomal or mitochondrial clusters (defined by Imaris segmentation) of AT2 cells are larger than those of AT1 and Cap2 cells (asterisk, p<0.0001). AT1 cells contain more lysosomal and mitochondrial clusters per cell than AT2 and Cap2 cells (asterisk, p<0.001). Lysosomes and mitochondria are closer to the nucleus in AT2 cells than in AT1 and Cap2 cells, attributable to cell size differences (asterisk, p<0.0001). GFP volume ration is calculated as the percentage of cell volume, approximated by GFP, occupied by lysosomes or mitochondria, and differs among cell types. Statistics is based on ordinary ANOVA with Tukey test.

### Live imaging of lysosome movement, fusion, and fission in embryonic lung epithelial progenitors

Expected as a major advantage of fluorescent fusion proteins, the multicolor kaleidoscope reporter in fresh tissues was sufficiently stable for simultaneous live-imaging of lysosomes, mitochondria, and microtubules, and even brighter than fixed tissues (Fig. 5, Supplementary Videos 3-6). Specifically, we activated the reporter with *Sox9^CreER^* in sparse lung epithelial progenitors and explant-cultured such labeled embryonic lungs. Over several hours, lysosomes and mitochondria mostly jittered but occasionally dashed across several microns, with a combined speed averaging 0.16 um/min and 0.14 um/min, respectively (Fig. 5A, Supplementary Video 4, 5). Moreover, the lysosomal puncta fused and fissioned, processes challenging to distinguish from restructuring for the mitochondrial network (Fig. 5A). EGFP-alpha-Tubulin was too diffuse under the current imaging condition to identify individual microtubule fibers, but reliably outlined cell nucleus, shape, and location (Supplementary Video 6).

**Figure 5:**
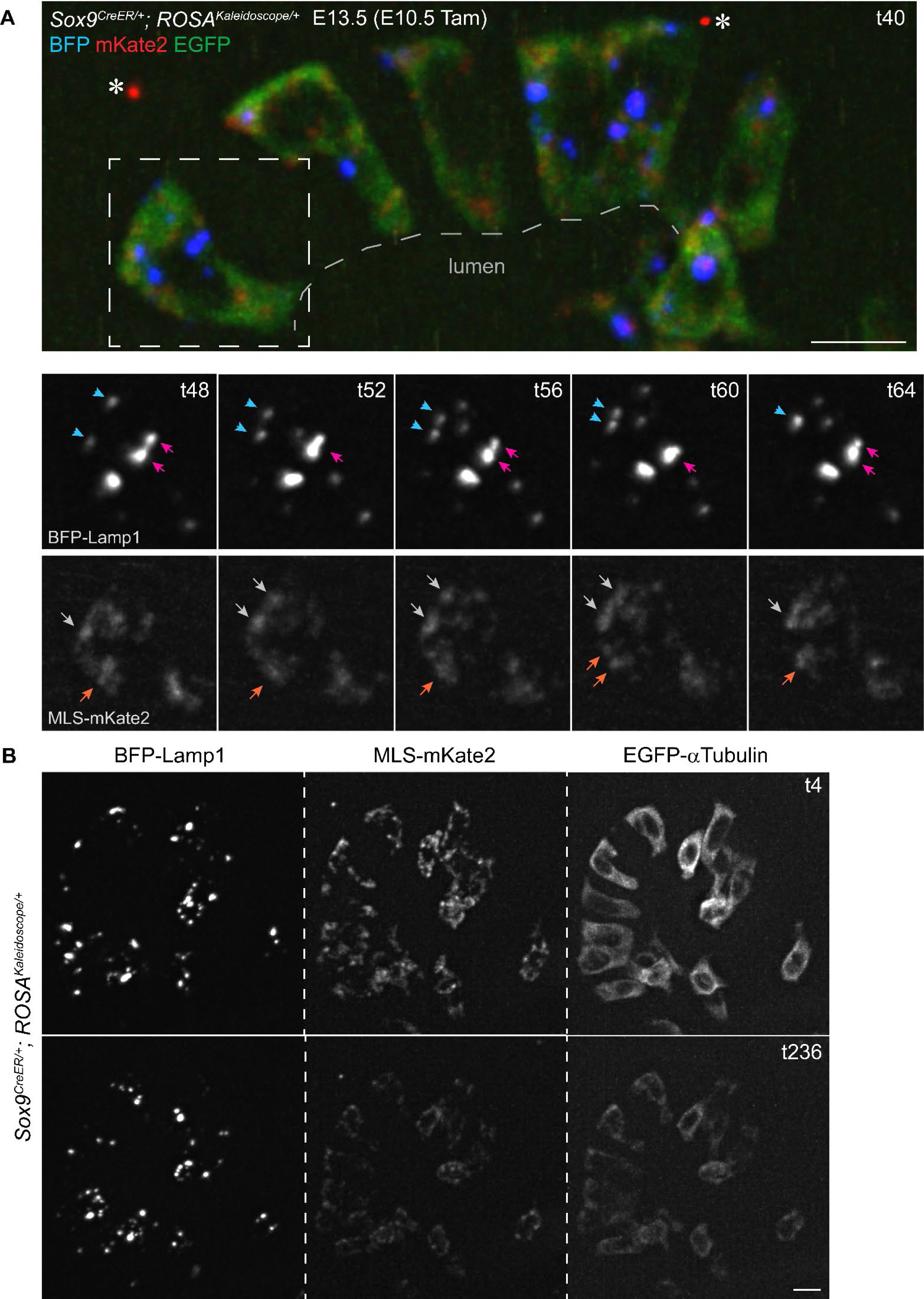
Live imaging of lysosomes and mitochondria in mouse embryonic lung epithelial cells in explant culture. (**A**) Native fluorescence from the 3 fusion proteins in *ROSA^Kaleidoscope^* activated in lung epithelial progenitors (*Sox9^CreER^*; Tam, 500 ug tamoxifen) at time 40 min (t40) in explant culture. Asterisk, artifact fluorescence. The boxed region is shown over time (t in minutes). Colored arrowheads track the same objects, consistent with lysosome fusion and fission as well as mitochondrial restructuring. Scale: 10 um. See also Supplementary Video 3-6. (**B**) Preservation of native fluorescence over 4 hr of live imaging. Lysosomal signal is more robust than mitochondrial and microtubule signals. Scale: 10 um.

### Organelle changes in AT1 cells after Sendai virus infection

After characterizing subcellular structures and dynamics in normal tissues, we applied the kaleidoscope reporter to a Sendai virus injury model, which features AT2-less regions where AT2 cells are depleted as a result of infection and AT1 cells temporarily seal the epithelial barrier before being displaced by invading airway basal-like cells^30^. Interestingly, unlike uninfected lungs and unaffected regions of infected lungs, AT1 cells in AT2-less regions had additional lysosomes but reduced mitochondria, subcellular changes preceding and potentially in preparation for their said displacement by increasing autophagy and reducing energy production (Fig. 6A, Fig. S4). Further supporting cell-type-specific organelle features, AT1 cells lineage-traced from AT2 cells were readily distinguishable from adjacent AT2 cells with their numerous dispersed lysosomes and mitochondria, predicting as drastic a change in subcellular structures as in gene expression during AT2-to-AT1 differentiation (Fig. 6B, Supplementary Video 7).

**Figure 6:**
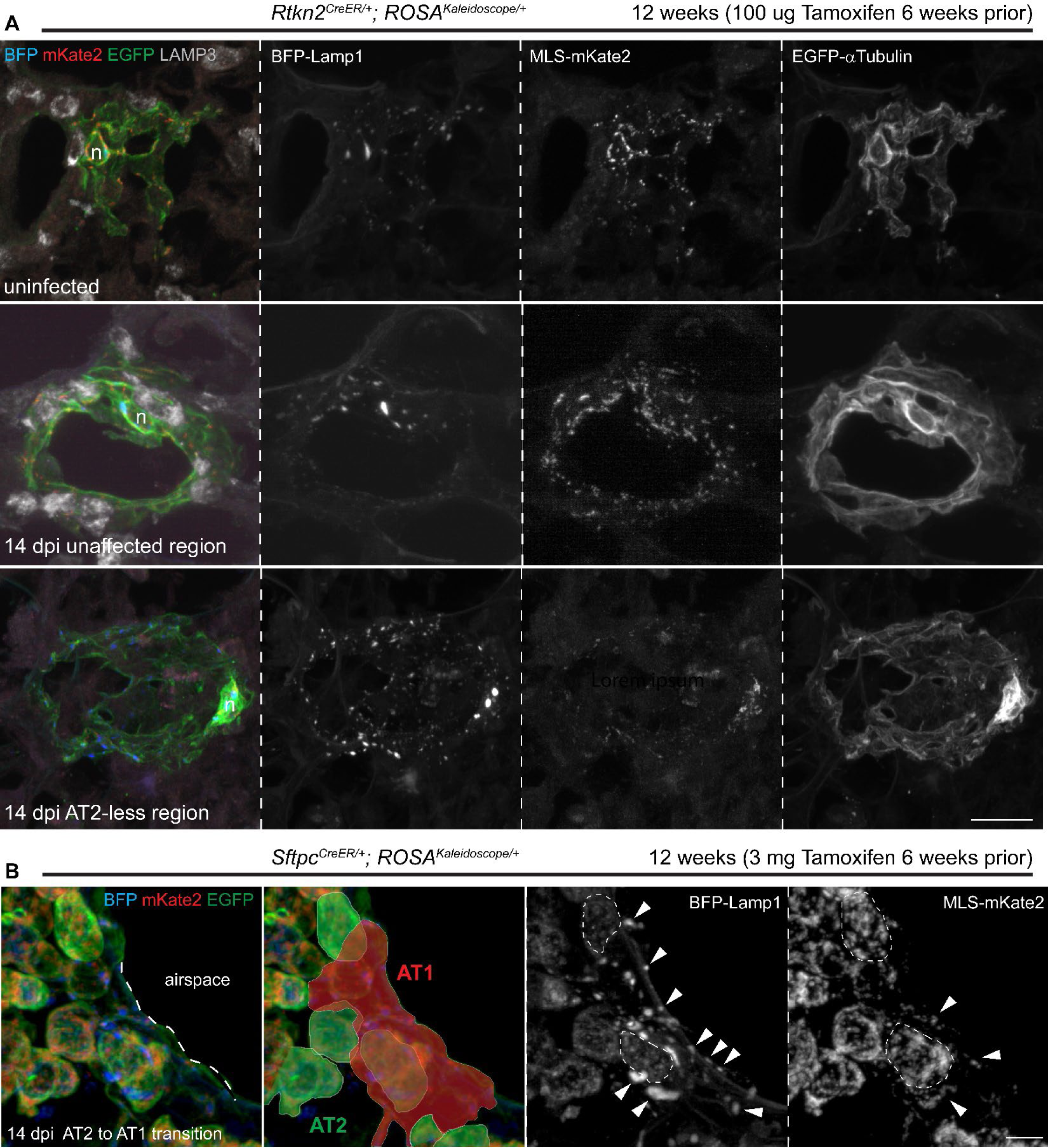
Lysosomal and mitochondrial changes in lung alveolar epithelial cells upon Sendai virus infection. (**A**) Genetically labeled individual AT1 cells using *Rtkn2^CreER^* in uninfected and 14 days post infection (dpi) with Sendai virus. Compared to uninfected lungs and unaffected regions in the infected lungs, AT1 cells in AT2-less regions, defined by lack of nearby LAMP3, have higher lysosomal fluorescence but lower mitochondrial fluorescence. Scale: 10 um. (**B**) Genetically labelled AT2 cells using *Sftpc*^*CreER*^ in lungs 14 days after Sendai virus infection. Compared to AT2 cells, a lineage-labeled AT1 cell (red mask) has characteristic numerous small lysosomes, supporting reorganization of subcellular structures as AT2 cells differentiate into AT1 cells. AT2 cells are outlined with dashes; AT1 cell organelles are marked by arrowheads. Scale: 10 um. See also Supplementary Video 7.

### *Yap/Taz* mutant lung epithelial cells accelerate lamellar body biogenesis

To further explore the notions that molecular events must be executed by subcellular machinery and that, in the same way that single-cell genomic tools capture molecular states, the kaleidoscope reporter captures subcellular states, we applied our reporter to a cellular analysis of the *Yap/Taz* mutant lung epithelial progenitors, which was shown via molecular analysis to undergo accelerated AT2 cell differentiation^29^. During normal AT2 cell differentiation, a AT2 cell-specific marker LAMP3 existed in two pools: multiple small puncta in progenitors and developing AT2 cells, and 1-2 large globules in mature AT2 cells, with the second pool overlapping with the kaleidoscope BFP-Lamp1 and gradually increasing in size (Fig. 7). Since large lysosomes were present in early progenitors even before the onset of LAMP3 expression and LAMP3-negative airway cells (Fig. 7A, 7B, S5A), LAMP3 was likely produced as small puncta that subsequently merged with lysosomes to form lamellar body clusters, consistent with them being lysosomal related organelles ^31, 32^. In the *Yap/Taz* mutant, both pools of LAMP3, distinguished by an overlap with BFP-Lamp1, were larger in size than those at the same and even later developmental stages (Fig. 7C). This enlargement in both vesicular and clustered LAMP3 could result from over-production of LAMP3 and related proteins due to transcriptional misregulation^29^, overwhelming subcellular machinery for lamellar body production, storage, and secretion. This lysosomal phenotype was consistent with the exaggerated AT2 cell differentiation predicted from molecular profiling^29^; by comparison, mutant cell mitochondria were not visibly different and mutant cell perimeters fell between AT1 and AT2 cells, possibly due to stretching to compensate for blockage of AT1 cell differentiation and expansion (Fig. S5B). These diverse cellular phenotypes, akin to differential and unaffected gene expression, highlighted the rich information herein and its use in defining biology.

**Figure 7:**
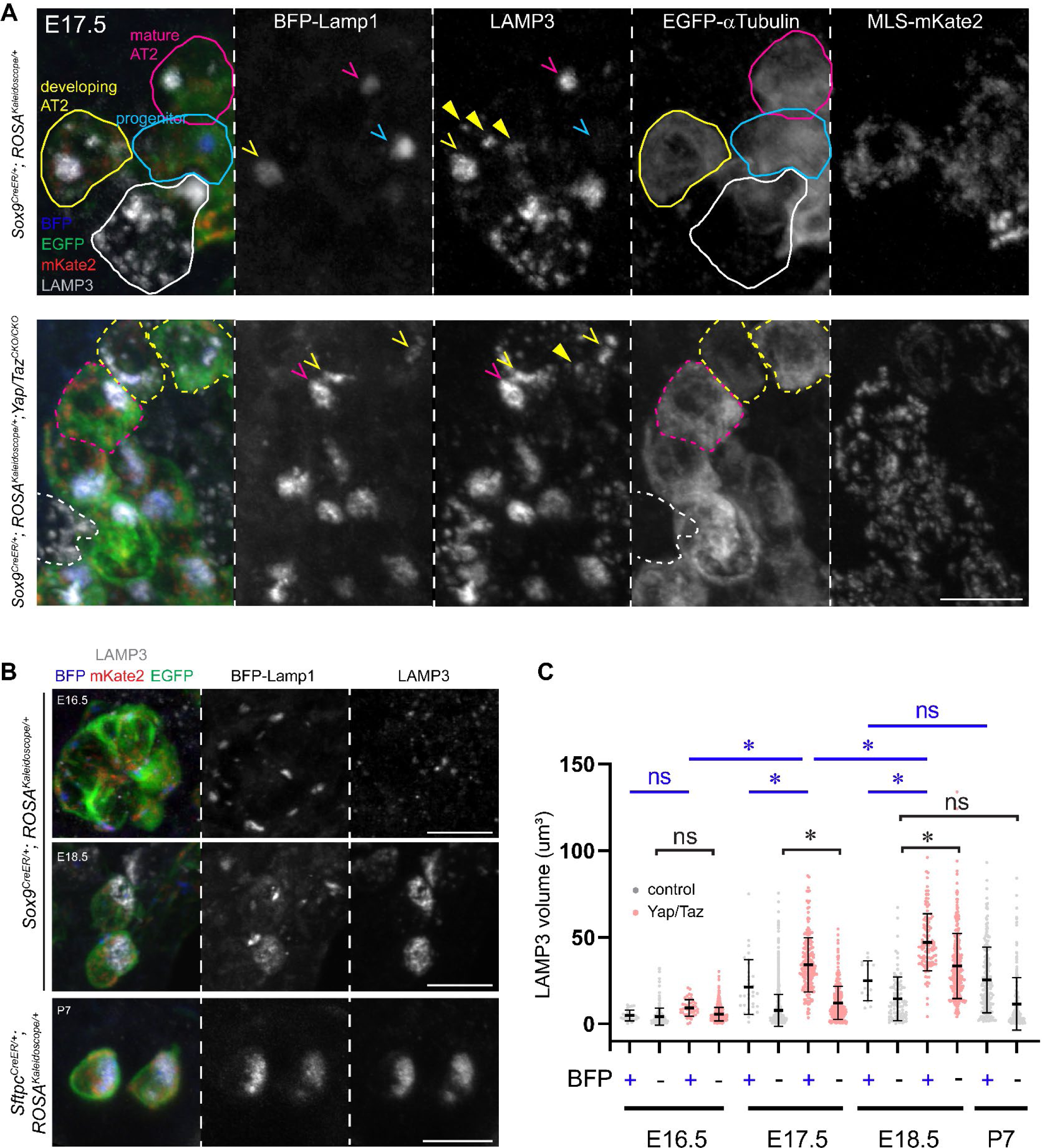
Premature lysosome maturation in *Yap/Taz* mutant cells. (**A**) LAMP3 particles shift from non-existent or small, scattered (arrowhead; BFP-) in progenitors and developing AT2 cells (blue and yellow, respectively) to large, overlapping with BFP-Lamp1 (caret; BFP+) in mature AT2 cells (magenta). Unrecombined cells have white outline. More AT2 cells are in the *Yap/Taz* mutant (dash) and at a more mature stage. 500 ug tamoxifen was given at E13.5 for the control; 2 mg tamoxifen was given at E14.5 for the *Yap/Taz* mutant. Scale: 10 um. (**B**) Normal temporal changes in lysosome size from progenitors at E16.5 (500 ug tamoxifen at E12.5) to AT2 cells at E18.5 (500 ug tamoxifen at E14.5) and P7 (300 ug tamoxifen at P1). Scale: 10 um. (**C**) Quantification of LAMP3 particle volumes over time grouped by overlapping with BFP-Lamp1, showing an earlier increase in volume and a larger end volume in the *Yap/Taz* mutant. Asterisk, p<0.05; ns, not significant (ordinary ANOVA with Tukey test; data are mean and standard deviation).

## DISCUSSION

This study represents an initial step to unravel cell-type-specific use of the subcellular machinery in tissues. We generate a Cre-dependent, tricolor fluorescent reporter mouse that simultaneously labels lysosomes, mitochondria, and microtubules across diverse cell types and with single-cell resolution. Applying this tool to the lung with entangled cellular processes of dozens of cell types, we document lysosomal and mitochondrial distributions in the convoluted AT1 and Cap2 cells and upon viral infection; track organelle dynamics in embryonic epithelial progenitors; and reveal accelerated lamellar body biogenesis in a *Yap/Taz* mutant.

Our study is motivated by the unmet need to extend the extensive knowledge of cell biology in cultured cells to that in native tissues. Such extension is necessary for the same reason that genes often function distinctly across cell types and in ways not recapitulated in culture. Notably, without native matrix, cellular, and geometric environments, cultured cells cannot mimic the full extent of cellular specialization needed for physiology, including endothelial cells in blood-filled vessels, folded ultrathin AT1 cells integrated with neighboring alveolar cells, and their aberrant versions from cumulative developmental deviation due to genetic mutations. Such cellular specialization is reflected in not only cell morphology, but also organelle usage such as mitochondrial subpopulations in skeletal muscle cells and lysosomal related organelles such as melanosomes and Weibel-Palade bodies^32, 33^. Using the tricolor reporter, we show that AT2 cell lamellar bodies, a type of lysosomal related organelles, form via gradual coalescence of LAMP3 vesicles with lysosomes and this process is accelerated upon *Yap/Taz* deletion (Fig. 7).

By establishing the feasibility of multicolor imaging of tissue cell biology, our study paves the way for systematic single-cell subcellular analysis, akin to single-cell genomics. First, leveraging fluorescent fusion proteins validated in cultured cells and other safe genomic loci such as *Hprt*, *H11*, *Tigre*, and *Col1a1*, the next generation of kaleidoscope mice would allow spectral imaging of all known subcellular structures including actin, intermediate filaments, Golgi, ER, endosomes, cell junctions, and matrix adhesions. The resulting kaleidoscopic cell-type-specific features represent a modern version of Golgi staining for cell morphology to classify neurons. Second, super-resolution imaging can be used to resolve microtubule dynamics as well as individual mitochondria and lysosomes versus their clusters and, more broadly, to map organelle interactomes in tissues such as mitophage^12^. Third, besides transcriptional regulators such as YAP/TAZ (Fig. 7), these subcellular reporters would enable real time mechanistic studies of direct regulators of cytoskeleton assembly and anchorage; vesicle budding, transport, and fusion; as well as cell junction establishment and remodeling.

Their Cre-dependent design will provide definitive evidence for intercellular transfer of organelles such as mitochondria ^34^. Last, therapeutic drugs targeting cell biology such as the microtubule stabilizer Taxol can be evaluated for side-effects on all cell types in tissues.

## MATERIALS AND METHODS

### Cloning

The ROSA targeting vector Ai9 was a gift from Hongkui Zeng (Addgene plasmid # 22799; http://n2t.net/addgene:22799; RRID: Addgene_22799). EGFP-Tubulin-6 was a gift from Michael Davidson (Addgene plasmid # 56450; http://n2t.net/addgene:56450; RRID: Addgene_56450). mTagBFP-Lysosomes-20 was a gift from Michael Davidson (Addgene plasmid # 55263; http://n2t.net/addgene:55263; RRID: Addgene_55263. The MLS-mKate2 plasmid was a generous gift from Dr. Anthony Barrasso and Dr. Ross Poche at Baylor College of Medicine.

The polycistronic cloning vector PTE2A-PGEMT^19^ was a generous gift from Dr. Li Qian at University of North Carolina. Fluorescent fusion protein sequences were amplified via high-fidelity PCR with custom primers to add cloning adapters (Table 1). The PCR fragments were digested to ligate into the PTE2A-PGEMT vector sequentially as in Figure 1S. The resulting PGEMT-BFP-Lamp1-P2A-MLS-mKate2-T2A-EGFP-Tubulin plasmid was sequence verified and digested with MluI (R0198, New England Biolabs) to replace the tdTomato sequence in Ai9. The resulting ROSA-Kaleidoscope plasmid was also treated with a Cre recombinase (M0298, NEB) for 30 min at 37°C to remove the transcriptional stop cassette to test the fusion proteins in culture.

**Table 1:**
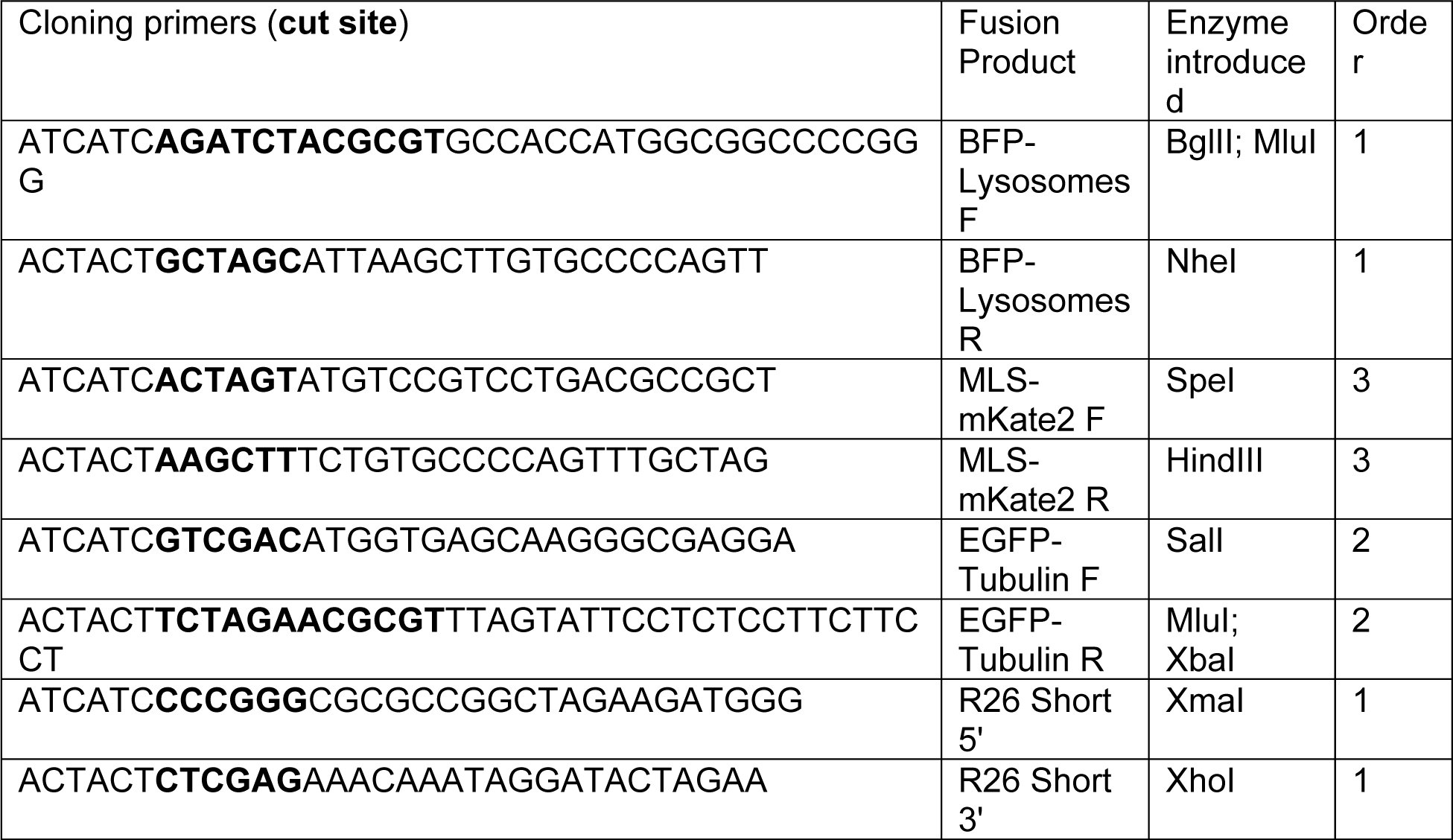
Cloning primers

### Mice

The following mouse lines were used: *Rtkn2^CreER^* ^29^, *S*ftpcCreER ^28^, *Sox9^CreER^* ^35^, CMV-Cre^24^, *Yap^CKO^* ^36^; *Taz^CKO^* ^36^.The *ROSA^Kaleidoscope^* knock-in allele was generated at the Genetically

Engineered Mouse Facility at MD Anderson Cancer Center using CRISPR targeting via standard mouse embryonic stem cells (mESCs) transfection. The Kaleidoscope targeting construct was digested with KpnI to linearize the DNA. Linear DNA was then diluted to 2.2 µg/µl and electroporated into 1 x 10^7^ G4 (hybrid 129/B6) mESCs. Transfected mESCs were selected for resistance to G418 and puromycin. DNA from individual surviving clones was screened by PCR analysis (see Table 2). Positive clones were then injected separately using standard procedures into blastocysts derived from albino-C57Bl/6N mice to generate male chimeras, which were then mated to black female C57Bl/6N to establish germline transmitted progeny.

**Table 2:**
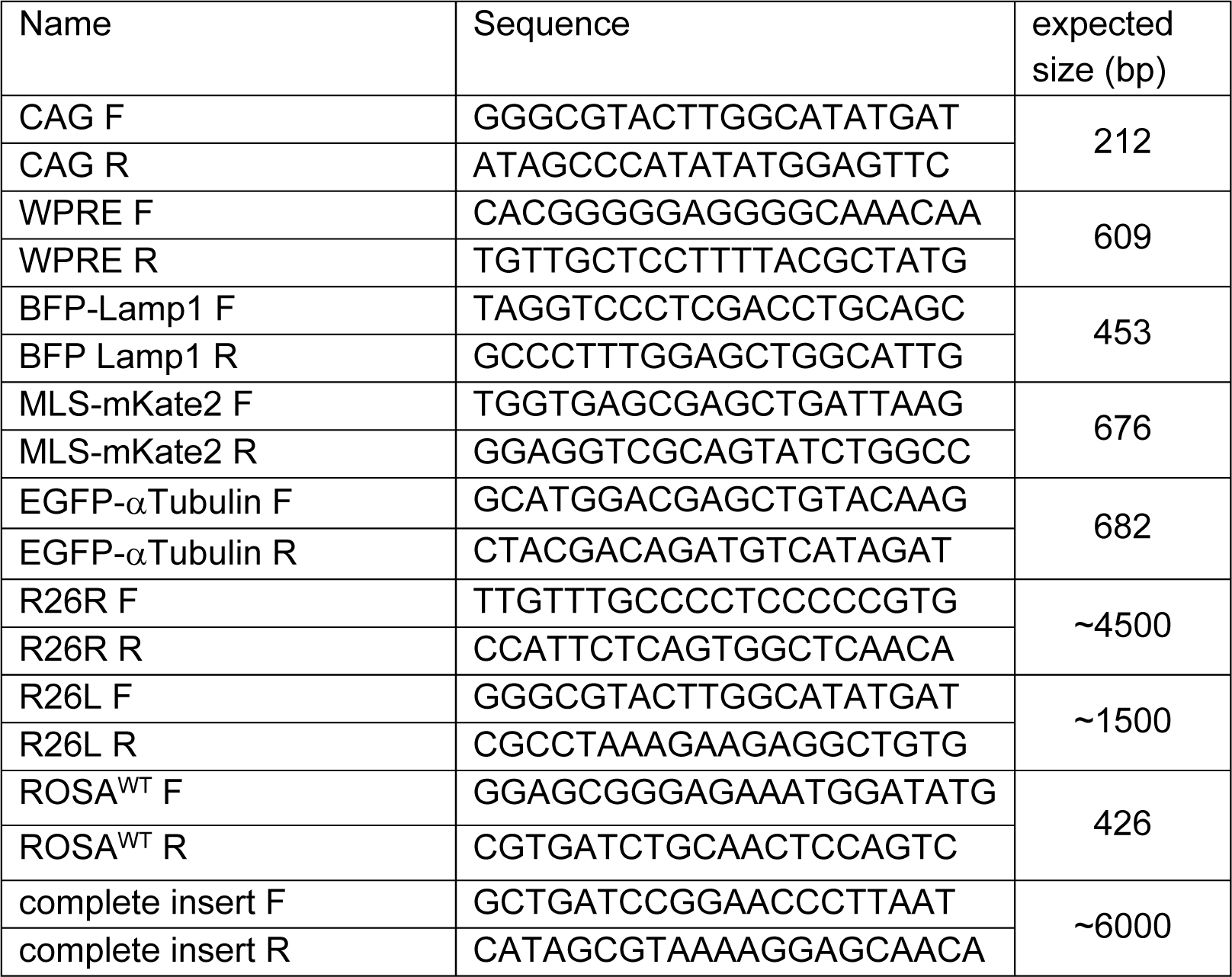
Genotyping primers and expected sizes

Presence of the targeted gene was established using PCR analysis of DNA obtained from tail samples (Table 2).

The *Car4^CreER^* allele was generated using CRISPR targeting via standard pronuclear injection^37^. Specifically, 400 nM gRNA (Synthego), 200 nM Cas9 protein (E120020-250ug, Sigma), and 500 nM circular donor plasmid were mixed in the injection buffer (10 mM Tris pH 7.5, 0.1 mM EDTA). The 5’ homology arm in the donor plasmid was PCR amplified between 5’- CCTAGAAACTCCCCAGGGATGC and 5’-TGGAGGCTTTATAGCAGAGCCACG; the 3’ homology arm was PCR amplified between 5’-GTGGCTTCATTGGTGATTGACCCT and 5’- CTTTAACATTATATTAGTTGCT. The gRNA targeted 5’-CCAGC<u>ATG</u>ATCGCGACGCCTAGG with the last 3 nucleotides being the protospacer adjacent motif (PAM; not included in gRNA) and a potential cryptic start codon in the 5’ UTR underlined and replaced by that of CreER used in Sox9*^CreER^* ^35^.

For embryonic experiments, pregnancies were designated as embryonic day (E) 0.5 on day of plug. Tamoxifen (T5649, Sigma) dissolved in corn oil (C8267, Sigma) was administered via intraperitoneal injection to pregnant mothers at doses and ages in figure legends to induce Cre recombination. Experiments were conducted in both male and female mice and investigators were aware of genotypes at the time of experiments. The mice were housed under conditions of 22°C, 45% humidity, and 12–12 h light–dark cycle. All animal experiments were approved by the Institutional Animal Care and Use at MD Anderson Cancer Center. Every effort was made to process samples in the same tubes as experimental and biological replicates to reduce variation between experiments.

### Cell line culture

HEK293T cells were grown in DMEM (D8418, Sigma) with 10% fetal bovine serum (FBS, 89510, VWR) and 1% PenStrep (15140, Gibco) on No.1 coverslips (48366-067, VWR) in 6-well plates (10062-892, VWR). After cotransfection using Lipofectamine 3000 (L3000-008, ThermoFisher) of ROSA-Kaleidoscope and pCAG-Cre (a gift from Connie Cepko; Addgene plasmid # 13775; http://n2t.net/addgene:13775; RRID:Addgene_13775), HEK293T cells were grown for 24-48 hr, fixed in 2% paraformaldehyde (PFA; P6148, Sigma), and immunostained or immediately imaged on an Olympus FV1000 confocal. MLE15 cells were grown in RPMI-1640 media (11875119, ThermoFisher) with 10% FBS, 1% PenStrep on No.1 coverslips in 6-well plates. Cells were transfected with Cre-recombined ROSA-Kaleidoscope using TransIT-2020 (MIR5404, Mirus Bio). For western blots, HEK293T cells were grown in 10 cm plates (25-202, GenClone) and transfected using lipofectamine 3000 with Cre-recombined ROSA- Kaleidoscope, BFP-Lamp1 (Addgene #55263), MLS-mKate2, or EGFP-alpha-Tubulin (Addgene #56450) and grown 24 hr. Cell lysis buffer (FNN0021, Invitrogen) was added to plates and centrifuged at 500 rpm for 3 min, followed by Pierce BCA analysis (#23227, ThermoFisher) on the supernatant.

### Primary cell culture

P7 CMV-Cre; *ROSA^Kaleidoscope/+^* lungs were removed after perfusion with phosphate buffered saline (PBS), freed of any surrounding tissue and trachea, and minced with forceps into ∼1 cm^3^ pieces. Tissue fragments were incubated in 1.35 mL PBS plus 150 uL 20 mg/mL Collagenase Type I (Worthington CLS-1 dissolved in PBS) and 15 uL 20 mg/ml DNase I (Worthington D dissolved in PBS) at 37°C for 40 min, with manual mixing halfway and at the end of incubation. Following digestion, tissue was washed with 10% FBS in Leibovitz media (21083-027, Gibco) and spun down. The supernatant was removed and cells incubated in red blood cell lysis buffer (15 mM NH_4_Cl, 12 mM NaHCO_3_, 0.1 uM EDTA, pH8.0) for 3 min before washing as before.

Cells were then resuspended in prewarmed RPMI-1640 with 10% FBS, 1% PenStrep and plated on No.1 cover slips in 6-well culture plates. Media was changed daily for 3 days to remove dead cells. On day 4, cells were fixed in 2% PFA for 2 hr, washed with PBS, mounted with Aquapolymount (18606, Polysciences) and imaged.

### Embryonic lung explant culture

Time-mated females were harvested at E14.5 and lungs with trachea and glottis were removed from embryos. The Cre-recombined ROSA-Kaleidoscope plasmid and Fast Green FCF dye (MKCD1540, Sigma) were mouth-pipetted through glottis until embryonic lumens were visibly filled. Lungs were electroporated in BioRad Xcell electroporation cuvettes (165-2088, BioRad) at 35 V for 3 rounds of 25 ms with 1s intervals. Lungs were then placed on Nuclepore Track-Etch Membrane filters (10417101, Whatman) and floated on RPMI-1640 media with 10% FBS 1% PenStrep for 24 hr before fixation in 0.5% PFA for 4 hr. After PBS wash, lungs were embedded in optimal cutting temperature compound (OCT; 4583, Tissue-Tek), cryosectioned, and imaged on Deltavision Deconvolution scope.

### Live imaging

Time-mated females were injected with 250 ug tamoxifen 2 days prior to sacrifice at E12.5 or E13.5. Embryos were screened for fluorescence and positive lungs were mounted on coverslip-bottom plates (P35G-1.5-10-C, Mattek) in a minimal amount of Leibovitz media. Lungs were imaged on a spinning disc confocal (3i Intelligent Imaging Innovations) using a 40x air objective at intervals indicated in figure legends at 37°C with 5% CO_2_ and humidified air. Following image acquisition, time-lapse videos were deconvolved using 3i software AutoQuant 3D Blind Deconvolution and converted to tiff files for import to Imaris software.

### Section immunofluorescence and confocal imaging

Postnatal and adult lungs were inflation-harvested as described (Yang et al., 2016) with minor modifications. Briefly, mice were sacrificed via intraperitoneal injection of avertin (T48402, Sigma) and perfused through the right ventricle with PBS. The trachea was cannulated and the lung inflated with 0.5% PFA in PBS at 25 cm H_2_O pressure, submersion fixed in 0.5% PFA at room temperature (RT) for 3-6 hr, and washed in PBS at 4°C overnight on a rocker. Section immunostaining was performed following published protocols with minor modifications (Alanis et al., 2014; Chang et al., 2013). Fixed lung lobes were cryoprotected in 20% sucrose in PBS containing 10% OCT at 4°C overnight and then embedded in OCT and frozen at -80°C. Sections were cut at 30 um thickness and blocked in 5% normal donkey serum (017-000-121, Jackson ImmunoResearch) in PBS with 0.3% Triton X-100 (PBST) before incubation with primary antibodies diluted in PBST in a humidified chamber at 4°C overnight. Sections were washed with PBS in a coplin jar for 30 min at RT then incubated with donkey secondary antibodies (Jackson ImmunoResearch) diluted in PBST at RT for 1 hr. After another 30 min wash with PBS, sections were mounted with Aquapolymount and imaged in ∼20 um z-stacks on a confocal microscope (FV1000V, Olympus) using a 60X oil objective. Liver, inner ear, and muscle tissues were removed following right ventricle perfusion and fixed in 0.5% PFA for 3-6 hr at RT and then cryoprotected and sectioned as above. See Table 3, 4 for a list of antibodies.

**Table 3:**
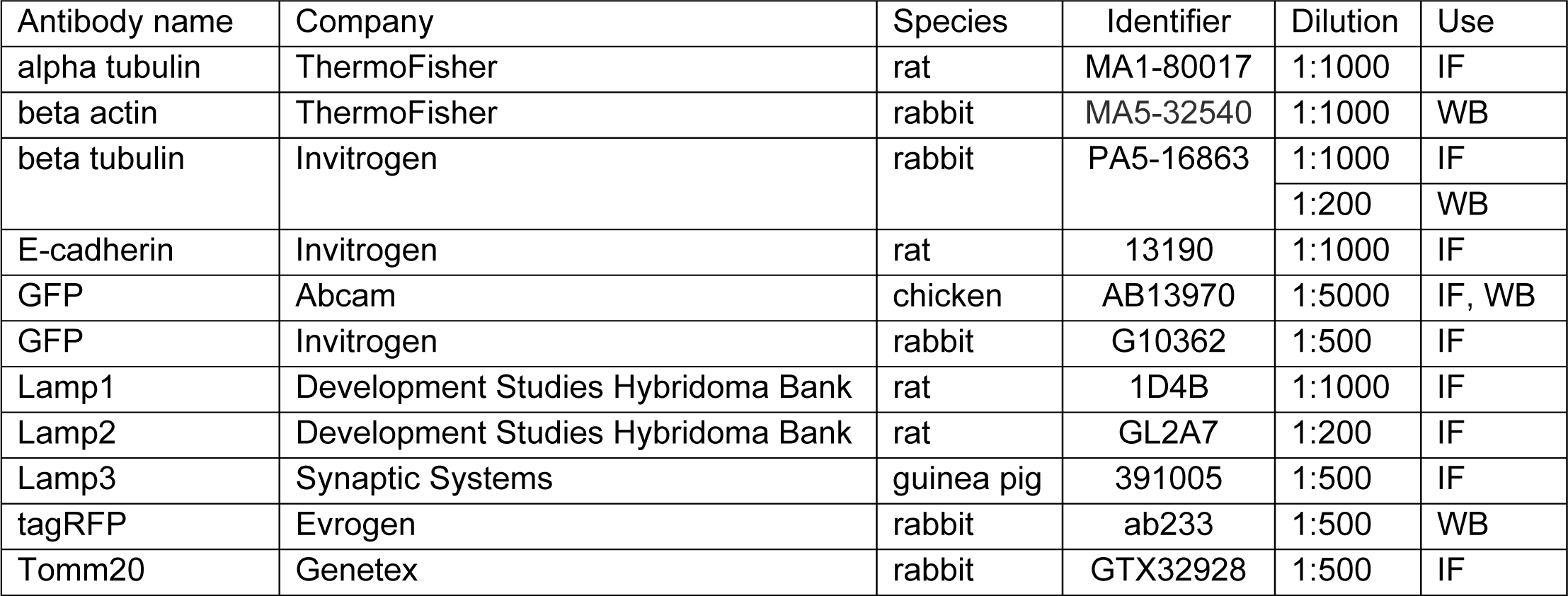
Primary antibodies and dilutions (IF, immunofluorescence; WB, western blot)

| Antibody name | Company | Species | Identifier | Dilution | Use |
| --- | --- | --- | --- | --- | --- |
| alpha tubulin | ThermoFisher | rat | MA1-80017 | 1:1000 | IF |
| beta actin | ThermoFisher | rabbit | MA5-32540 | 1:1000 | WB |
| beta tubulin | Invitrogen | rabbit | PA5-16863 | 1:1000 | IF |
|  |  |  |  | 1:200 | WB |
| E-cadherin | Invitrogen | rat | 13190 | 1:1000 | IF |
| GFP | Abcam | chicken | AB13970 | 1:5000 | IF, WB |
| GFP | Invitrogen | rabbit | G10362 | 1:500 | IF |
| Lamp1 | Development Studies Hybridoma Bank | rat | 1D4B | 1:1000 | IF |
| Lamp2 | Development Studies Hybridoma Bank | rat | GL2A7 | 1:200 | IF |
| Lamp3 | Synaptic Systems | guinea pig | 391005 | 1:500 | IF |
| tagRFP | Evrogen | rabbit | ab233 | 1:500 | WB |
| Tomm20 | Genetex | rabbit | GTX32928 | 1:500 | IF |

### Whole mount immunostaining

This was performed following published protocols with minor modifications (Yang et al., 2016). In brief, ∼3 mm wide strips from the edge of the left lobes of postnatal or adult lungs or whole lobes of embryonic lungs were blocked with 5% normal donkey serum in PBST for 2-4 hr at RT and then incubated with primary antibodies diluted in PBST overnight at 4°C on a rocker. The strips were then washed with PBS+1%Triton X-100+1%Tween-20 (PBSTT) on a rocker at RT 3 times over 4 hr. Secondary antibodies diluted in PBST were incubated on a rocker overnight at 4°C. On the third day, the strips were washed as above before fixation with 2% PFA in PBS for at least 2 hr on a RT rocker. Finally, the strips were mounted on slides using Aquapolymount with the flattest side facing the coverslip. Z-stack images of 20-40 um thick at 1 um step size were taken from the top of the tissue to obtain an en face view.

### Western blot

CMV-Cre; *ROSA^Kaleidoscope/+^* postnatal lungs were removed after PBS perfusion and flash frozen. Tissue was placed in cell lysis buffer (FNN0021, Invitrogen) and protease inhibitor cocktail (78442, Thermo Scientific) and mechanically agitated twice for 45 seconds with chrome steel beads (11079113c, BioSpec Products). Following 10 min incubation on ice, samples were centrifuged at 13,000 rpm for 15 min and supernatant retained. Pierce BCA analysis was performed to determine protein content (23227, ThermoFisher). 40 ug protein was run on a variable percentage acrylamide gel (4561094, BioRad) for 1 hr at 115 V before transferring to protein membrane (IPV00010, Millipore) at 50 V for 2 hr. Membranes were blocked in 5% bovine serum albumin (BSA, A3059, Sigma) in tris-buffered saline (A3059, Sigma) with 0.1% Tween-20 (TBST pH 7.4) at 4°C for 1 hr with constant agitation before incubation with primary antibodies (Table 3) overnight at 4°C with constant agitation. 3-5 short washes in TBST were performed before incubation in 2% BSA in TBST with relevant secondary antibodies (Table 4). Chemiluminescence was imaged on BioRad ChemiDoc gel imager with ∼800 ul Immobilon Crescendo Western HRP substrate (WBLUR0100, Millipore). Blots were treated with Restore™ Western Blot Stripping Buffer (21059, ThermoFisher) for 30 min at RT with agitation before blocking and staining as before, up to 2 additional times.

**Table 4:**
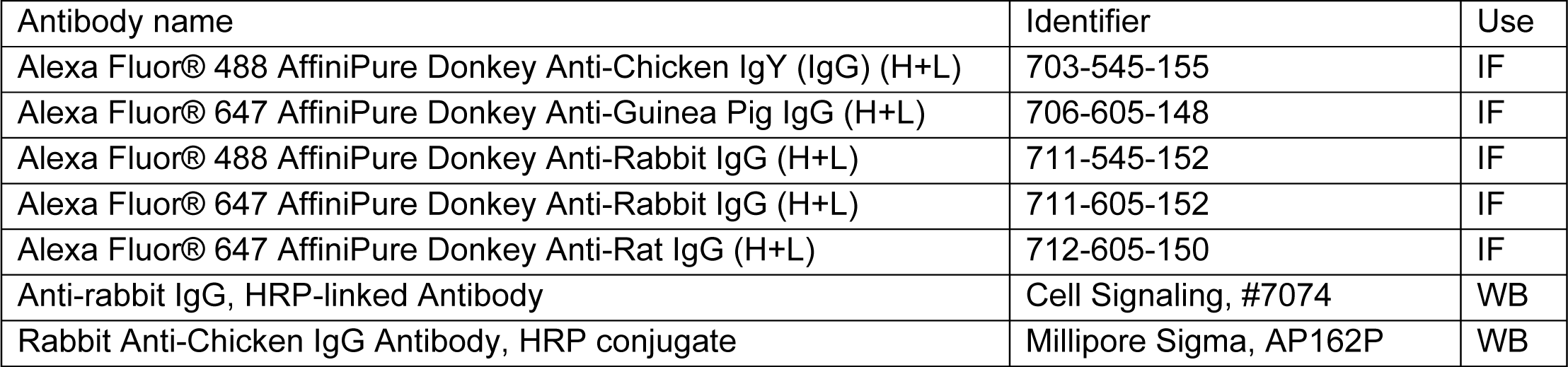
Secondary antibodies (IF, immunofluorescence; WB, western blot) All IF secondary antibodies sourced from Jackson ImmunoResearch and used at 1:1000. All WB antibodies were used at 1:5000 dilution

| Antibody name | Identifier | Use |
| --- | --- | --- |
| Alexa Fluor® 488 AffiniPure Donkey Anti-Chicken IgY (IgG) (H+L) | 703-545-155 | IF |
| Alexa Fluor® 647 AffiniPure Donkey Anti-Guinea Pig IgG (H+L) | 706-605-148 | IF |
| Alexa Fluor® 488 AffiniPure Donkey Anti-Rabbit IgG (H+L) | 711-545-152 | IF |
| Alexa Fluor® 647 AffiniPure Donkey Anti-Rabbit IgG (H+L) | 711-605-152 | IF |
| Alexa Fluor® 647 AffiniPure Donkey Anti-Rat IgG (H+L) | 712-605-150 | IF |
| Anti-rabbit IgG, HRP-linked Antibody | Cell Signaling, #7074 | WB |
| Rabbit Anti-Chicken IgG Antibody, HRP conjugate | Millipore Sigma, AP162P | WB |

### Sendai virus infection

Viral infection was performed as described^30^. Briefly, isoflurane-anesthetized mice were suspended by the upper incisors and treated with a non-lethal dose (∼2.1 X10^7^ plaque-forming units) Sendai Virus (ATCC #VR-105, RRID:SCR_001672CSCSSCS) in 40 ul PBS or PBS only via oropharyngeal instillation. Lungs were inflation harvested at 14 days post infection, fixed, cryoprotected, and sectioned as described above.

### Image analysis

Imaris 9.9 (Bitplane) was used for image analysis. Surfaces were created using DAPI (BFP- Lamp1), 488 (EGFP-aTubulin), 555 (mito-mKate2) and 647 (antibody stains) for volume, distance, size, and speed measurements. Identical surface creation parameters were used between experiments. Manual editing for aberrant surfaces was performed prior to statistical analysis.

## ACKNOWLEDGEMENTS

We thank the University of Texas MD Anderson Genetically Engineered Mouse Facility for generating the *ROSA^Kaleidoscope^* and *Car4^CreER^* mice, with the support of the Cancer Center Support Grant (CA #16672). We thank Dr. Adriana Paulucci-Holthauzen at the Microscopy Laboratory of the University of Texas MD Anderson Center for guidance on microscopy acquisition and image processing, with the support of the NIH instrumentation grant (S10OD024976). We thank Drs. Hongkui Zeng, Michael Davidson Anthony Barrasso, Ross Poche, and Li Qian for sharing plasmids. We thank Dr. Harold Chapman for providing the *S*ftpcCreER mice. This work was supported by the University of Texas MD Anderson Cancer Center Retention Fund, and National Institutes of Health R01HL130129 and R01HL153511 (JC).

## AUTHOR CONTRIBUTIONS

VH and JC designed research; VH, AL, and AMGG performed research and analyzed data; JC generated the *Car4^CreER^* mice; VH and JC wrote the paper; all authors read and approved the paper.

## COMPETING INTERESTS

The authors declare no competing interests.

## Supplementary Figures

**SupFig. 1:**
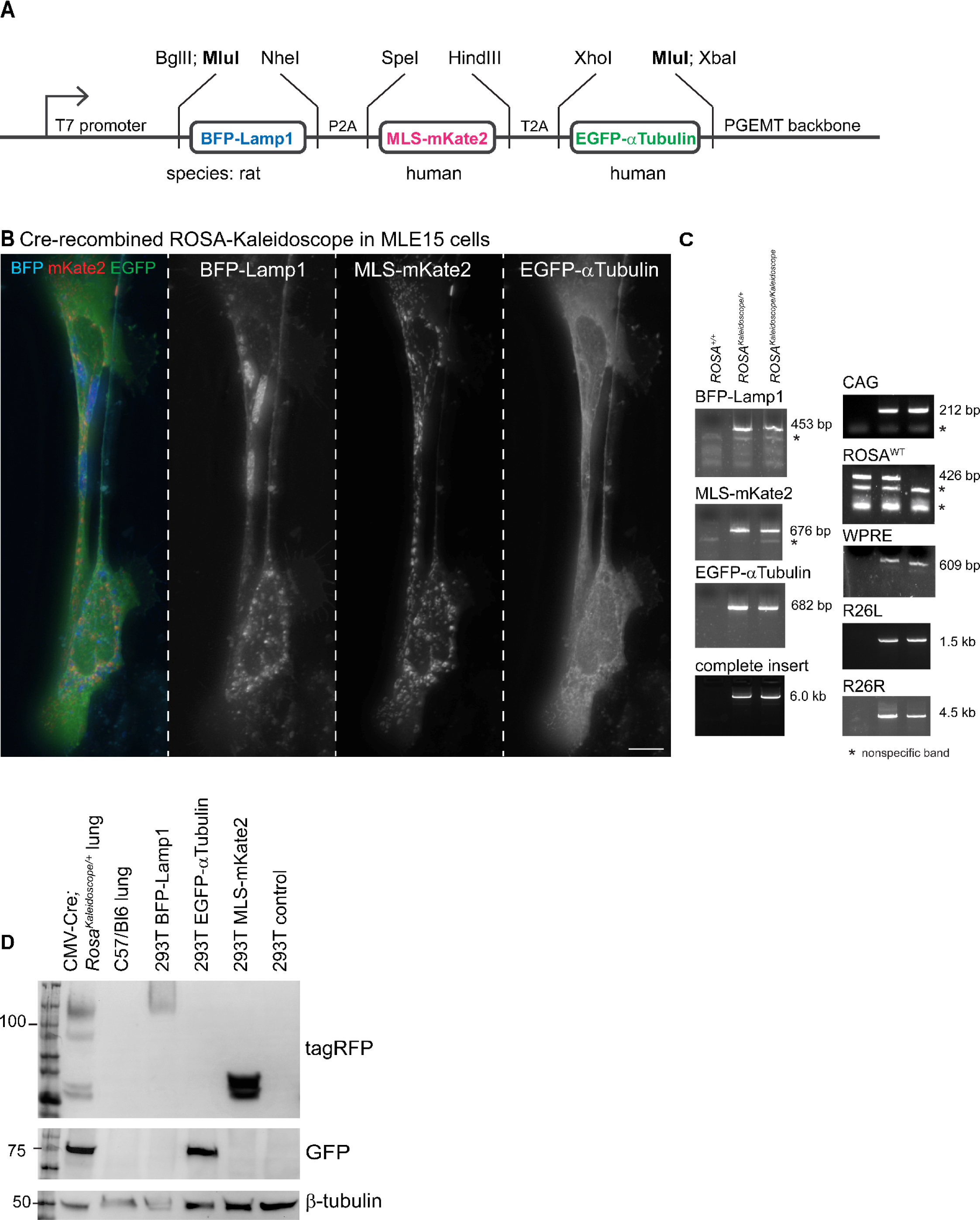
Additional details on the design and validation of the Kaleidoscope reporter. (A) Intermediate vector to show cloning sites to concatenate the 3 fusion proteins. The bolded MluI sites used to transfer them to the Ai9 targeting vector to generate the ROSA-Kaleidoscope plasmid. The Lamp1 sequence is from rat, while the MLS and alpha-Tubulin sequences are from human. See Table 1 for cloning primer sequences and the order of cloning. (B) MLE15 cells transfected with Cre-recombined ROSA-Kaleidoscope to show comparable fluorescence patterns as in HEK 293T cells. Scale: 10 um. (C) Expected PCR genotyping results of *ROSA^Kaleidoscope^* littermates. See Table 2 for genotyping primer sequences. (D) Western blots of lungs expressing all 3 fusion proteins and HEK 293T cells expressing individual fusion proteins. The second largest band specific to the Kaleidoscope lungs might be due to incomplete glycosylation of LAMP1. Both BFP and mKate2 are tagRFP derivatives and detected by the same antibody.

**SupFig. 2:**
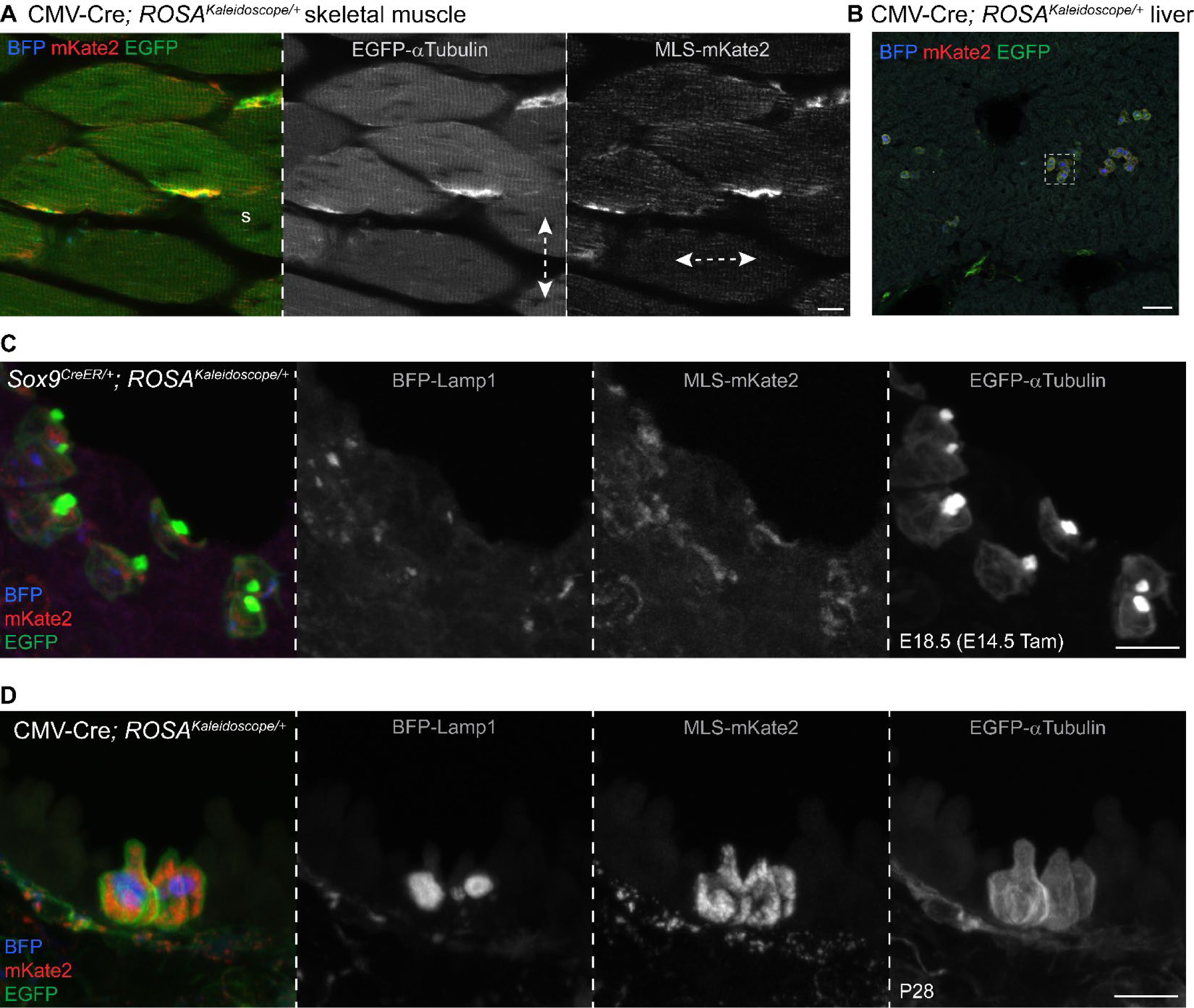
Additional characterization of the Kaleidoscope reporter in organs. (A) A single optical section of the skeleton muscle in Fig. 2B to show that microtubules are aligned perpendicular to actomyosin bundles, whereas mitochondria are along and between actomyosin bundles (double arrow). s, satellite cell. Scale: 10 um. (B) A low magnification view of the liver showing inefficient recombination by CMV-Cre. Boxed area is shown in Fig. 2B. Scale: 50 um. (C) Distribution of Kaleidoscope fusion proteins in airway ciliated cells with characteristic apical microtubule clusters. Tam, 500 ug tamoxifen. Scale: 10 um. (D) Distribution of Kaleidoscope fusion proteins in dome-shaped airway club cells and spindle-shaped airway smooth muscle cells. Scale: 10 um.

**SupFig. 3:**
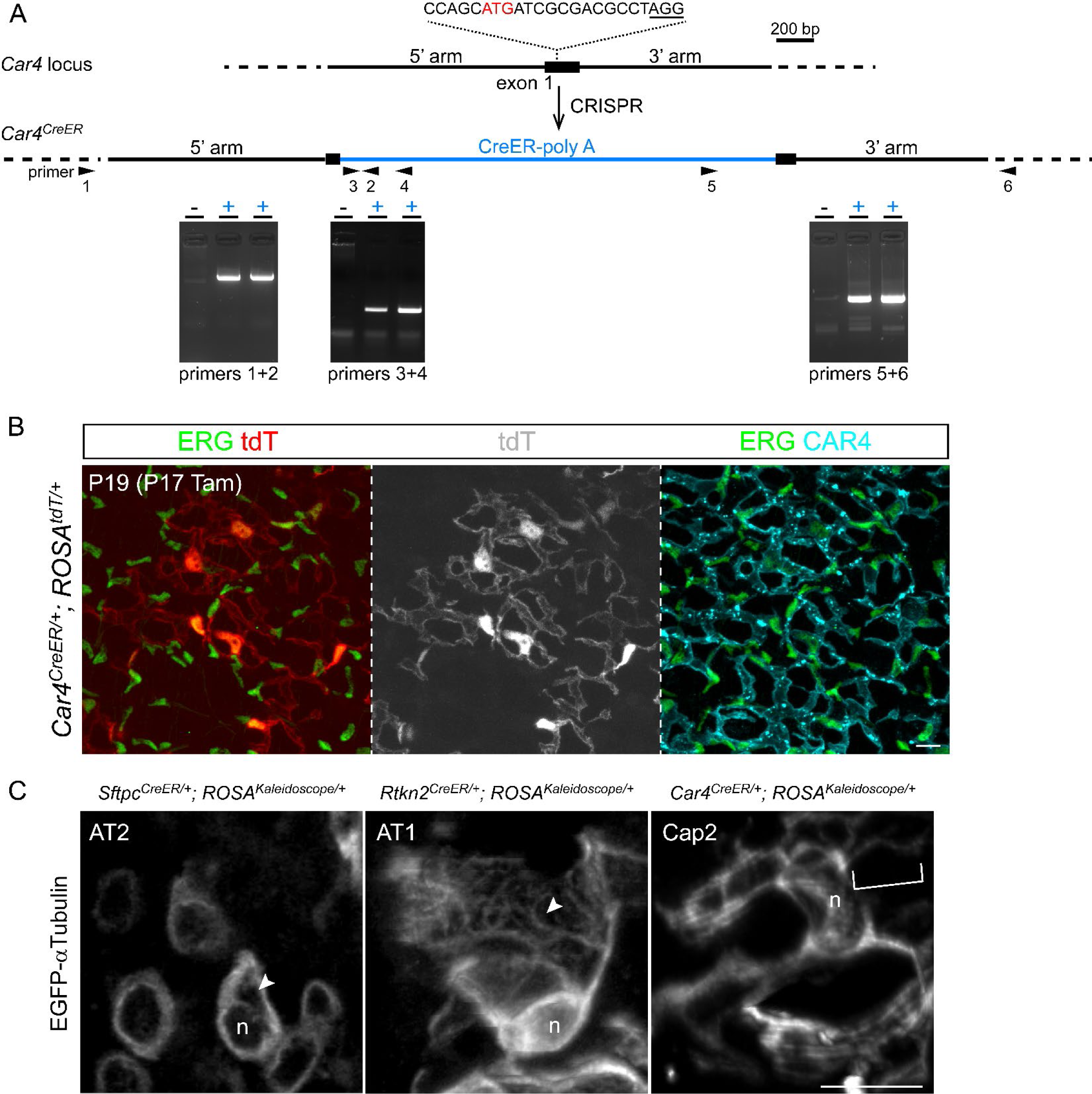
Generation of *Car4^CreER^* mice and high-magnification views of microtubules in lung cells. (A) CRISPR targeting of the *Car4* locus. The gRNA sequence is shown with the protospacer adjacent motif (PAM; not included in gRNA) underlined; a cryptic translation start codon of *Car4* (ATG in red) is replaced by that of CreER. Locus-specific PCR genotyping identifies two positive mice. The primer sequences are: #1, 5’-TGTACTGCTATTCCTTGTTCATCT; #2, 5’-AGAAGCATTTTCCAGGTATG; #3, 5’-TGACACTGAGAACCACAAACGGC; #4, 5’-GTTCGCAAGAACCTGATGGACA; #5, 5’-TTCTAGTTGTGGTTTGTCCAAACT; #6, 5’- AACGTGAACAGCTACAAGGCACTG. (B) *Car4^CreER^* targets Cap2 endothelial cells expressing ERG and CAR4 with characteristic large net-like morphology. Further characterization will be described in another study. Scale: 10 um. (C) Filamentous microtubules are apparent in thin z stacks (<5 um) of labelled AT2, AT1, and Cap2 cells (arrowhead and bracket). n, nucleus. See Fig. 4 for experimental conditions. Scale: 10 um.

**SupFig. 4:**
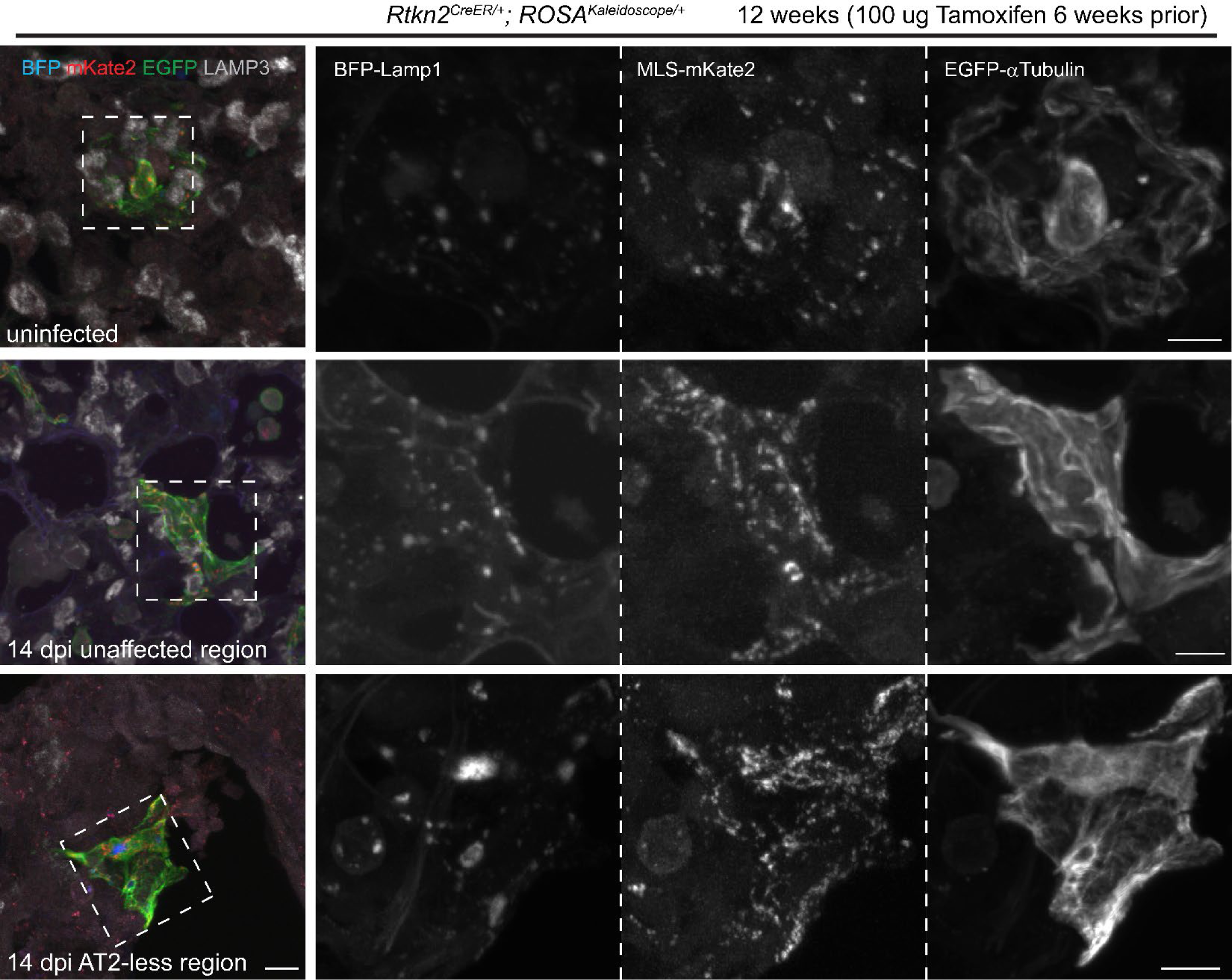
Additional examples of labelled AT1 cells upon Sendai virus infection. Low-magnification view of individual labeled AT1 cells in uninfected lungs and unaffected and AT2-less regions (devoid of LAMP3+ AT2 cells). Boxed regions are magnified. Mitochondrial fluorescence of the AT1 cell in the AT2-less region is not as reduced as in Fig. 5 possibly reflecting an earlier stage of cell displacement. Scale: 10 um.

**SupFig. 5:**
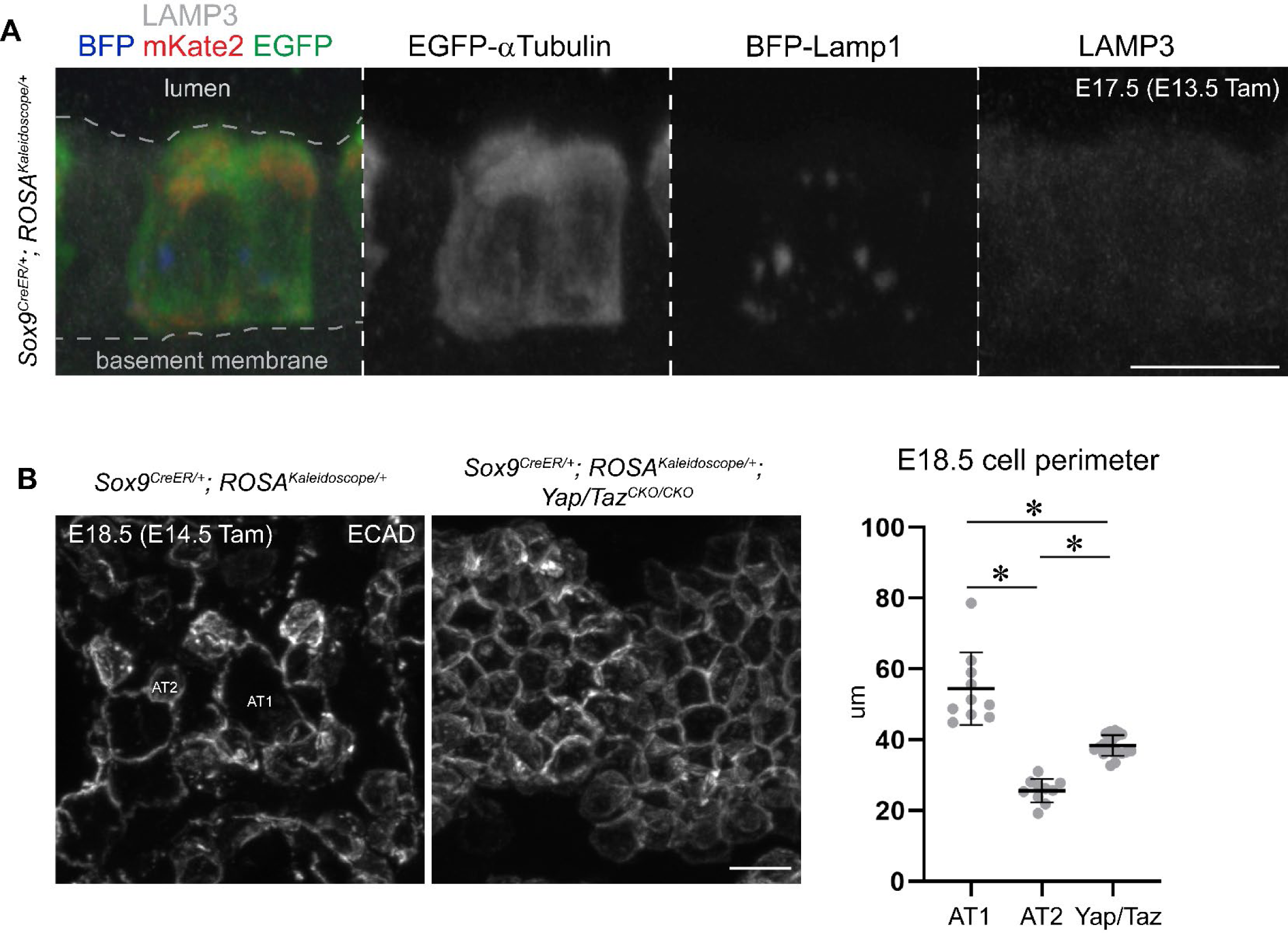
Further characterization of labeled epithelial cells in the *Yap/Taz* mutant (A) Labeled LAMP3- airway cells still contain BFP-Lamp1 puncta. Tam, 500 ug tamoxifen. Scale: 10 um. (B) En face view and perimeter quantification of alveolar epithelial cells outlined with E-Cadherin (ECAD) immunostaining. Cells in the *Yap/Taz* mutant have an intermediate perimeter between AT1 and AT2 cells of the control lung. Tam, 3 mg tamoxifen. Asterisk, p<0.0001 (ordinary ANOVA with Tukey test; data are mean and standard deviation). Scale: 10 um.

**SupVideo 1: 3D view of labeled AT1 cells, as in** Fig. 4A.

**SupVideo 2: 3D view of labeled Cap2 cells, as in** Fig. 4A.

**SupVideo 3: A 4-hr movie (4 min/frame) of live imaging of Sox9^CreER^ labeled epithelial cells in explanted embryonic lungs, as in** Fig. 5A**, showing all 3 fusion proteins.**

**SupVideo 4: A 4-hr movie (4 min/frame) of live imaging of Sox9^CreER^ labeled epithelial cells in explanted embryonic lungs, as in** Fig. 5A**, showing BFP-Lamp1.**

**SupVideo 5: A 4-hr movie (4 min/frame) of live imaging of Sox9^CreER^ labeled epithelial cells in explanted embryonic lungs, as in** Fig. 5A**, showing MLS-mKate2.**

**SupVideo 6: A 4-hr movie (4 min/frame) of live imaging of Sox9^CreER^ labeled epithelial cells in explanted embryonic lungs, as in** Fig. 5A**, showing EGFP-alpha-Tubulin.**

**SupVideo 7: 3D view of *Sftpc*^*CreER*^ lineage-labeled AT1 and AT2 cells, as in** Fig. 6B.

## Notes

### Competing Interest Statement

The authors have declared no competing interest.

